# Neural mechanisms underlying the precision of visual working memory

**DOI:** 10.1101/330118

**Authors:** Yijie Zhao, Shuguang Kuai, Theodore P. Zanto, Yixuan Ku

## Abstract

The neural mechanisms associated with the limited capacity of working memory has long been studied, but it is still unclear how the brain maintains the fidelity of representations in working memory. Here, an orientation recall task for estimating the precision of visual working memory was performed both inside and outside an fMRI scanner. Results showed that the trial-by-trial recall error (in radians) was correlated with delay period activity in the lateral occipital complex (LOC) during working memory maintenance, regardless of the memory load. Moreover, delay activity in LOC also correlated with the individual participant’s precision of working memory from a separate behavioral experiment held two weeks prior. Furthermore, a region within the prefrontal cortex, the inferior frontal junction (IFJ), exhibited greater functional connectivity with LOC when the working memory load increased. Together, our findings provide unique evidence that the LOC supports visual working memory precision, while communication between the IFJ and LOC varys with visual working memory load.

## Introduction

Working memory (WM), a system that maintains and manipulates information in a short period for goal-directed actions (Baddeley 2012), is a critical cognitive function supporting everyday behaviors including language comprehension (Baddeley 2003), learning (Mayer and Moreno 1998) and reasoning (Conway et al. 2003; Luck and Vogel 2013) and correlated with general intelligence (Engle et al. 2011). However, the capacity of visual WM (VWM) is severely limited (Luck and Vogel 2013) and likely related to posterior brain activity (Todd and Marois 2004, 2005; Vogel and Machizawa 2004). Recent studies have further shown that the precision of VWM representation is also restricted (Bays and Husain 2008; Zhang and Luck 2008; Bays et al. 2009), while the neural mechanisms underlying such limited precision are still in debate.

Some studies have suggested elevated activity in the primary sensory cortex during WM maintenance may reflect WM representations, at least in the somatosensory domain (Zhou and Fuster 1996, 2000). However, other research have reported an absence of persistent activity in early sensory regions during VWM maintenance (Luna et al. 2005; Offen et al. 2009), Despite a lack of elevated activity during WM maintenance, detailed visual features such as orientation (Ester et al. 2009; Harrison and Tong 2009), motion direction (Emrich et al. 2013) and spatial location (Sprague et al. 2014, 2016) can be decoded in human early visual cortices by multivariate analysis of neuroimaging data (Riggall and Postle 2012; Albers et al. 2013). These recent findings imply the precision of VWM is encoded in early visual cortices. Yet, activities in the superior intraparietal sulcus (IPS) and the lateral occipital complex (LOC) depend on object complexity during the delay (Xu and Chun 2006), which suggests additional neural regions may contribute to the precision of VWM. Therefore, additional research is required to understand the roles of these regions in VWM precision.

Persistent neuronal activity in the prefrontal cortex has long been associated with the maintenance of VWM contents when visual stimuli are no longer present (Fuster and Alexander 1971; Goldman-Rakic 1995). Moreover, top-down signals from the prefrontal cortex modulate activity within sensory cortices and influence WM processes (Gazzaley and Nobre 2012). Transcranial magnetic stimulation (TMS) to the prefrontal cortex alters sensory processes in visual cortex during both visual perception (Ruff et al. 2006) and VWM (Zanto et al. 2011) tasks. Importantly, this top-down modulation of visual cortical activity during sensory encoding correlates with alterations in VWM performance (Bollinger et al. 2010; Zanto et al. 2011). However, whether this top-down modulation persists during the delay, and whether such modulation during VWM maintenance relates to VWM precision, still remains unknown. Together, both sensory cortex and prefrontal cortex have been identified as important regions supporting VWM, but their process-specific contributions remain unclear. Indeed, fronto-parietal cortices and sensory cortices have been proposed to represent different aspects of WM, such as the quantity (i.e., capacity) and the quality (i.e., precision) of WM representations, respectively (Ku et al. 2015), but support for this hypothsis is lacking.

In order to elucidate neural networks associated with VWM quantity and quality, we asked subjects to perform an orientation recall task inside a magnetic resonance imaging (MRI) scanner. A whole brain univariate analysis was applied to explore the neural candidates associated with the precision of VWM, as univariate methods have less bias towards sensory regions than multivariate methods when decoding visual features (Jimura and Poldrack 2012; Davis et al. 2014). We further assessed the connectivity between the precision-sensitive areas and other regions that may support VWM precision processes.

## Materials and methods

### Experiment 1: Behavioral experiments

#### Participants

Twenty-six healthy right-handed volunteers (10 males, age range 21.31±1.94) from the East China Normal University participated in the behavioral experiment. The experimental protocol was approved by the Ethics Committee of the School of Psychology and Cognitive Science, East China Normal University. Informed written consents were obtained from all subjects.

### Stimuli and Procedure

Stimuli were presented on a 60 Hz LCD monitor through MATLAB-based Psychophysics Toolbox (Version 3).

Each trial began with a fixation cross presented in the center of the screen throughout the whole trial. Subjects were asked to fixate on the cross (Fig. 1) during the experiment. A 200-ms sample array consisting of one, two, three, four or six bars were then presented on the screen. All items were located on invisible circle with a radius of 4^°^. Each item in the sample array was 1.5^°^ × of visual angle. The orientations of the bars were chosen from 1^°^ to 180^°^. In order to prevent between-item distraction, orientations of every two bars were differentiated at least 15 ^°^. After the sample array, one 900-ms retention interval with a blank screen was presented. In the probe array, one of the bars was presented in the same spatial location, but with a random orientation, which required the subjects to recall the orientation of the corresponding item from the sample array by rotating the bar with the computer mouse. Subjects were required to perform it as precisely as they can without a time limitation. Inter-trial-interval (ITI) was jittered between 200 to 500 ms with a 50 ms step. Every subject completed five blocks, each consisted of 100 trials (20 trials for each WM load) and the sequence of WM load was randomly ordered in each block.

**Figure 1.**
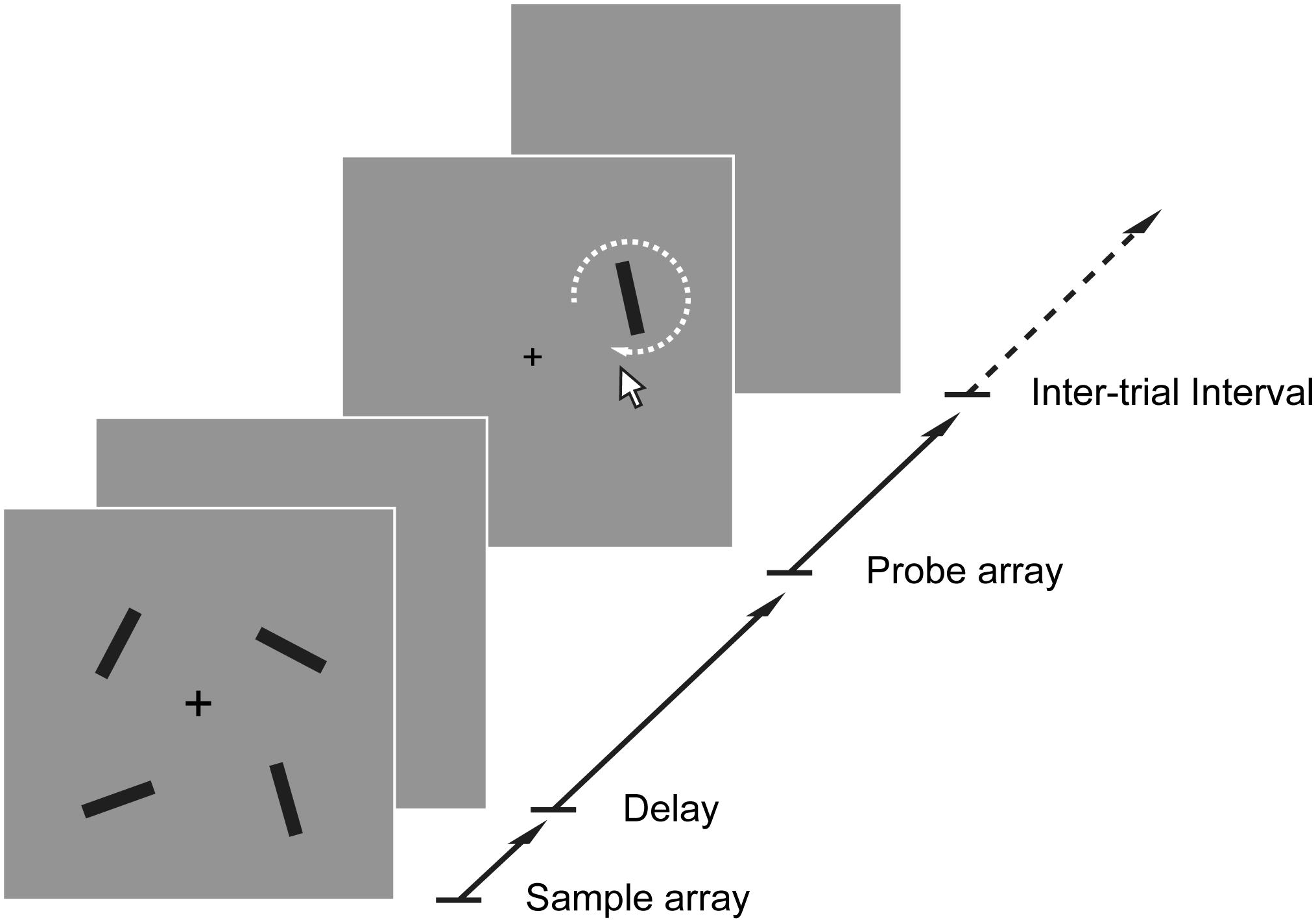
Orientation recall task with an example of WM load 4. On each trial, subjects were shown an arrangement of several bars presented at 4^±^ eccentricity. After a delay period, one of the bars was presented again and subjects were asked to adjust the bar using a computer mouse to match the orientation of it in their memory. In the behavioral experiment, the sample array and delay array lasted 0.2 s and 0.9 s, respectively, and the probe array lasted until the subject response. In the fMRI experiment, the sample, delay, and probe array lasted 0.5 s, 8 s, and 4 s, respectively. Inter-trial intervals were chosen randomly from 0.2 to 0.5 s (with 50 ms steps) in the behavioral experiment and 3.5 s, 5.5 s or 7.5 s in the fMRI experiment.

#### Behavioral Data Analyses

The data from each subject contained a set of the distance between the reported orientation and the origin one, which reflected the response error ranging -π to π. The standard mixture model (Zhang and Luck 2008; Bays et al. 2009) was used to fit the performance of each subject in each memory load condition according to a Maximum likelihood estimation. There were three possible sources in the model: a uniform distribution for the trials, which was not encoded into memory, a Von Mises distribution for the trials that the target orientations were encoded into memory and another Von Mises distribution with the same concentration as the first one but centered at nontarget responses. Correspondently, four parameters were returned from the model while the summary of the first three parameters equaled one: Pt, the probability that the target item was remembered; Pn, the probability that nontarget items were remembered by mistake; Pu, the guess rate and SD, the standard deviation of the Von Mises distribution, which reflected the precision of the memory representation. This model can be described as the following equation:

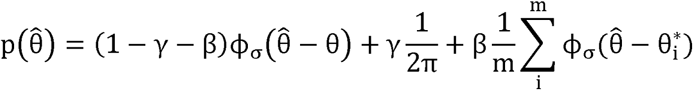

where θ is the origin orientation (in radians), 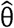 is the reported orientation, γ refers to the proportion of trials that subjects reported randomly, α is the probability of successfully reporting the target orientation, β is the probability of reporting a nontarget orientation and Φ _σ_ is the Von Mises Distribution with mean zero and standard deviation σ.

For more details of this model, see Bays et al. (2009) (Bays et al. 2009). Parameters were estimated using MATLAB toolbox available at http://www.bayslab.com and separate estimations were obtained for each subject and condition.

### Experiment 2: fMRI experiment

#### Participants

Fifteen subjects from experiment 1 volunteered to take part in the fMRI experiment. Two subjects were excluded because of excessive head-movement (> 3 mm), leaving thirteen subjects (8 males, age range 21.54±1.99) for data analysis. The two experiments were separated over at least two weeks to avoid practice effects. Subjects all completed Informed Written Consent and the experimental protocol was also approved by the Ethics Committee of the School of Psychology and Cognitive Science, East China Normal University.

### Stimuli and Procedure

The stimuli and procedure in the fMRI experiment (Fig. 1) were the same as the behavioral experiment except for: a. the sample array was presented for 500 ms; b. the duration of the delay period was 8 sec; c. there was a 4-s time limitation to respond to the probe (no subjects failed to respond within this time period in any trials); d. the ITI was extended to 3.5 s, 5.5 s or 7.5 s pseudorandomly; e. the memory load was 1, 2, 4 or 6 in this experiment.

Subjects performed 160 trials over eight runs in the scanner. Each run contained sixteen WM trials with four trials in each WM load and four 16-s blank trials with only the fixation cross presented on the screen. The probability of memory load switch was balanced. Stimuli were presented by a 60 Hz projector and viewed through a coil-mounted mirror.

### fMRI Data Acquisition

All MRI images were collected at the Shanghai Key Laboratory of Magnetic Resonance of the East China Normal University using a 3T Siemens Trio and a 12-channel RF coil. T1-weighted anatomical images were acquired using MPRAGE sequence (TR = 2530 ms, TE = 2.45 ms, flip angle: 7 ^°^, FOV: 256 mm, voxel size: 1 mm × 1 mm × 1 mm). For functional data, 33 slices covering the whole brain were defined: slice thickness 4 mm, slice gap 0 mm, FOV: 210 mm, phase-encode direction anterior-posterior. A T2^*^-weighted, gradient-echo EPI sequence was used: matrix size: 64 × 64, TR = 2 s, TE = 30 ms, flip angle: 90^°^, voxel size: 3.3 mm × 3.3 mm × 4 mm.

### fMRI Data Analyses

Functional data were pre-processed using the SPM8 (Wellcome Trust Centre for Neuroimaging, London, UK; www.fil.ion.ucl.ac.uk) toolbox in MATLAB. All volumes performed slice time correction, motion correction using the rigid-body to align all volumes to the first volume in the first run, co-registration with the subject-native anatomical volume, normalization to the Montreal Neurological Institute (MNI) space, spatial smoothing using an 8-mm FWHM Gaussian filter and 1/128 Hz cutoff high-pass temporal filter.

In our first general linear model (GLM), we seek to identify brain areas which show a main effect of WM load. According to previous theories, there exists a limitation of WM capacity which is the “magical number 4” (Cowan 2010). So we separated all trials into two categories: low WM load (combination of loads 1 and 2) and high WM load (combination of loads 4 and 6). Then, we modeled the sample array, delay and probe array including two levels of memory loads as separated regressors, producing a total of 6 regressors in the GLM. Regressors were modeled as stick function convolved with a standard hemodynamic response function (HRF). Six rigid body parameters were also included to correct for the head motion artifact. For the group level analysis, the “high WM” and “low WM” contrasts corresponding to the delay stage were subjected to second-level random effects analysis using a paired t-test (“high WM > low WM”) in SPM8.

In order to identify brain areas associated with VWM precision, we did another GLM analysis in which the absolute value of response errors (in radian) of every trial inside the scanner were put into the GLM as a parametric regressor. In this case, we pooled all response errors together regardless of the load condition. At the group level, we used a one-sample t-test to assess the brain areas that showed a negative association between the behavioral performance (i.e., less error means higher precision) and BOLD signal. Brain areas identified in this analysis were used as the Regions of Interest for WM precision (“precision ROIs”).

To investigate the functional connectivity between sensory areas and other parts of the brain during the delay period of VWM, a psychophysiological interaction (PPI) (Friston et al. 1997) analysis was performed. Volumes of interest (VOIs) were defined as 5 mm-radius sphere centered around areas showing the WM precision effect. The eigenvariate of the time series from these VOIs (or seeds) were extracted based on the first GLM. A PPI contrast of “high load > low load” was defined. At the group level, a one-sample t-test was applied to search for brain areas that show higher functional connectivity with these seeds in high load than in low load conditions.

All analyses were set to a threshold of p < 0.001 at the voxel level with a 70-voxel cluster extent to achieve a family-wise error corrected p < 0.05 using alpha probability simulations in the REST toolbox (http://www.restfmri.net (Song et al. 2011)).

### Within-subject Correlation Between Experiments

The activity in LOC during the delay period was related to the behavioral performance trial-by-trial inside the scanner. Furthermore, we tested whether the averaged activity in LOC inside the scanner predicted the precision of VWM performance outside the scanner. Here we applied a within-subject correlation approach (Bland and Altman 1995; Emrich et al. 2013) to examine the relationship between *changes* in the LOC response and *changes* in VWM precision across load. Specifically, we made the BOLD signal in LOC as the outcome variable and the behavioral performance of VWM precision as predictor variables, and we treated the subject variable as the covariate. In other words, an ANCOVA was used to eliminate between-subject differences of BOLD signal in LOC and then we can get the variation of LOC activity across WM loads due to changes of VWM precision estimated from the behavioral experiment outside the MRI scanner using the mixture model.

## Results

### Experiment 1: Behavioral experiment

In order to evaluate visual working memory precision for line orientation, subjects were asked to finish a 45-minute behavioral experiment (Fig.1). As showed in Figure 2, the height of the response error distribution curves declined and the width of the curves increased with an increasing WM load. Repeated measures ANOVA results revealed significant load effects on the probability of correct target response (Pt, F_(4,100)_ = 42.562, *p* < 0.001, 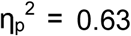), memory precision (F_(4,100)_ = 79.811, *p* < 0.001, 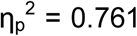), as well as on the probability of mistakenly reporting a nontarget item (Pn, F_(4,100)_ = 13.304, *p* < 0.001,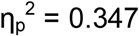) and the guessing rate (Pu, F_(4,100)_ = 10.373, *p* < 0.001, 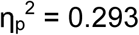).

**Figure 2.**
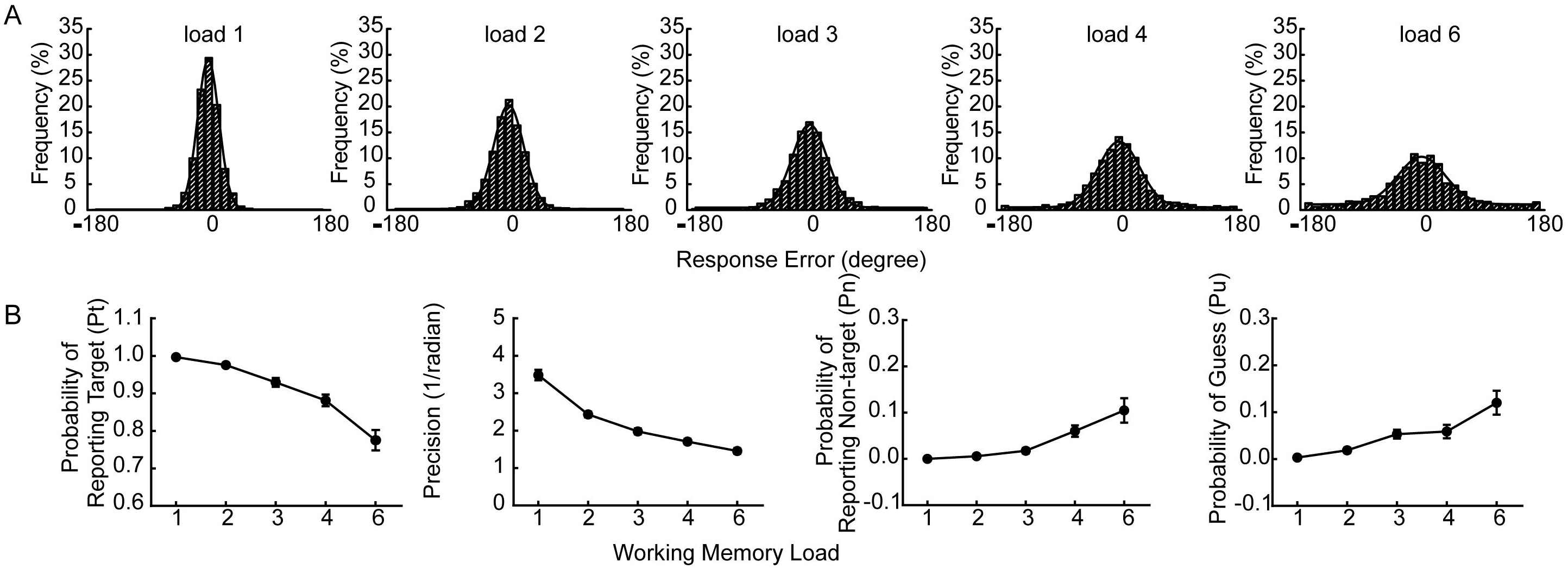
Behavioral experiment results. A. The probability distribution of the difference between reported orientation and original orientation (response error) across all subjects, along with the fit of a mixture model (solid line, see Methods) (Bays et al. 2009). The height of the distribution represents the probability of remembering the probed item while the width of the distribution quantifies the precision of working memory. B. Parameters of the mixture model as a function of load size. Error bars represent standard error (s.e.m.) across subjects. Results showed a decrease in memory capacity (Pt) and precision as a function of memory load, as well as an increase in non-target responses (Pn) and guessing rate (Pu) at higher memory loads.

### Experiment 2: fMRI experiment

#### Behavioral results

The behavioral results replicated the load effects from the behavioral experiment, such that a repeated measures ANOVA yielded the same main effects for load: Pt (F_(3,36)_ = 30.695, *p* < 0.001,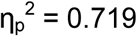), precision, (F_(3,36)_ = 104.593, *p* < 0.001,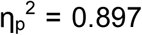), Pn (F_(3,36)_= 6.015, *p* = 0.002,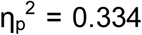), and Pu (F_(3,36)_ = 7.244, *p* < 0.001,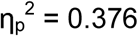).

## fMRI results

We built our first general linear model (GLM) to identify brain areas showing a WM load effect. Consistent with previous studies (Todd and Marois 2004; Barch et al. 2013), group level contrasts between “high WM load” and “low WM load” (i.e., the load effect) evoked widely spread activities in a frontal-parietal network (Fig. 3) during the delay period of the WM task. Specifically, seven cortical regions showed significant activations after multiple-comparison correction: bilateral frontal eye fields (FEF), bilateral inferior frontal junction (IFJ), dorsal anterior cingulate cortex (ACC), and bilateral inferior parietal lobe (IPL) (see Table 1).

**Table 1.**
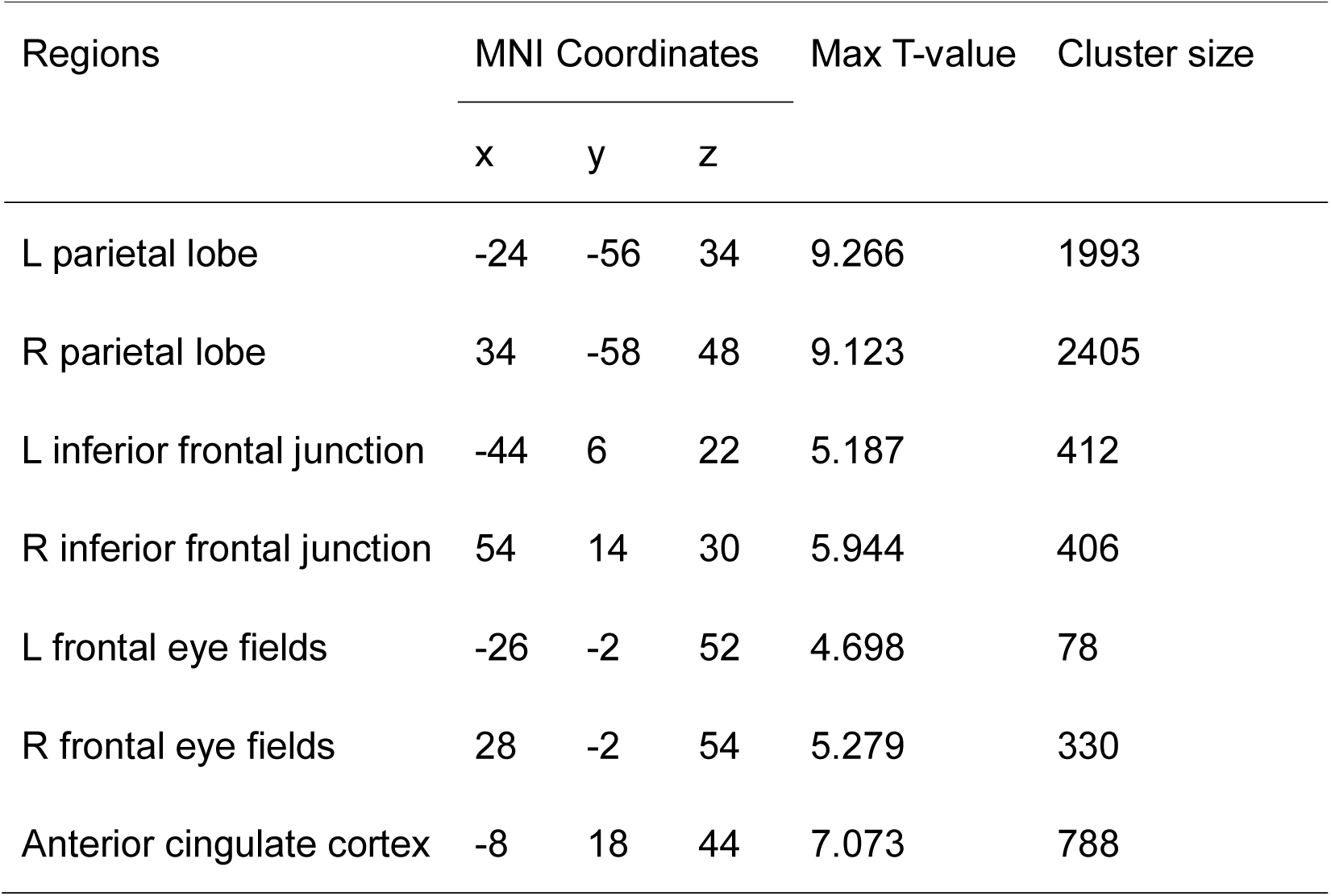
Results of whole-brain univariate analysis of the load effect

| Regions | MNI Coordinates |  |  | Max T-value | Cluster size |
| --- | --- | --- | --- | --- | --- |
|  | x | y | z |  |  |
| L parietal lobe | -24 | -56 | 34 | 9.266 | 1993 |
| R parietal lobe | 34 | -58 | 48 | 9.123 | 2405 |
| L inferior frontal junction | -44 | 6 | 22 | 5.187 | 412 |
| R inferior frontal junction | 54 | 14 | 30 | 5.944 | 406 |
| L frontal eye fields | -26 | -2 | 52 | 4.698 | 78 |
| R frontal eye fields | 28 | -2 | 54 | 5.279 | 330 |
| Anterior cingulate cortex | -8 | 18 | 44 | 7.073 | 788 |

**Figure 3.**
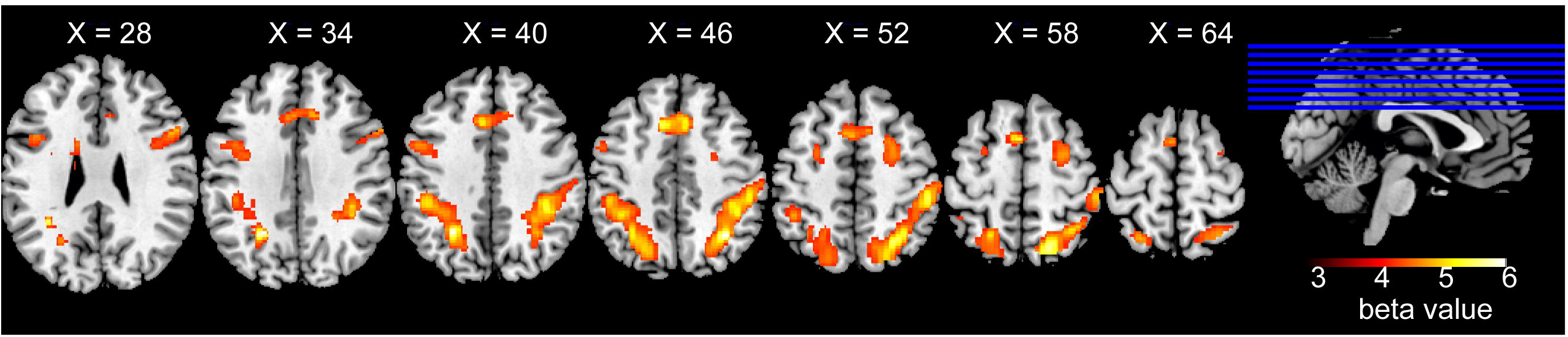
Activation related to working memory load (i.e., high WM load > low WM load) during the delay period. All data were analyzed on the MNI template. Results showed that voxels in bilateral parietal lobes, inferior frontal gyrus, medial frontal gyrus and anterior cingulate cortex (see Table 1) had significant activation differences between high / low WM load.

To investigate the neural representation of VWM precision, a parametric analysis was applied by putting the response errors (in radians) of each trial as a parametric regressor into our second GLM in which we pooled all response errors together regardless of the load condition. Results revealed that only three clusters of voxels in the lateral occipital complex (LOC) were negatively related with the increase of response errors in each trial during the delay period (Fig. 4A): two in the left LOC (peak MNI coordinate: [−26 −88 6], maximum t_(12)_ = 4.915, *p*_FWE_ < 0.05, 88 voxels; peak MNI coordinate: [−36 −80 −2], maximum t_(12)_ = 5.388, *p*_FWE_ < 0.05, 120 voxels), and one in the right LOC (peak MNI coordinate: [32 −80 2], maximum t_(12)_ = 6.058, *p*_FWE_ < 0.05, 73 voxels). Given the interpretation that the BOLD signal was larger in these three areas when behavioral error was smaller, the results indicated that these regions of interest (ROIs) may play an important role in holding the precise orientations in VWM, and serve as an candidate for “precision regions-of-interest (ROIs)”. Nevertheless, according to the results of experiment 1 and previous findings (Zhang and Luck 2008; Bays et al. 2009; van den Berg et al. 2012), VWM precision decreases with increasing memory load. So, it is possible that the regions tracing the precision might just reflect a load effect. Although we didn’t find any activation in the visual cortex when we directly measured the load effect using paired t-tests, it was still worth testing this possibility by adding WM load (low *vs.* high) as a regressor into the second GLM. No regions showed differential activation between the two levels of WM load, which confirmed the results that the three lateral occipital areas were only sensitive for memory precision rather than the memory load.

**Figure 4.**
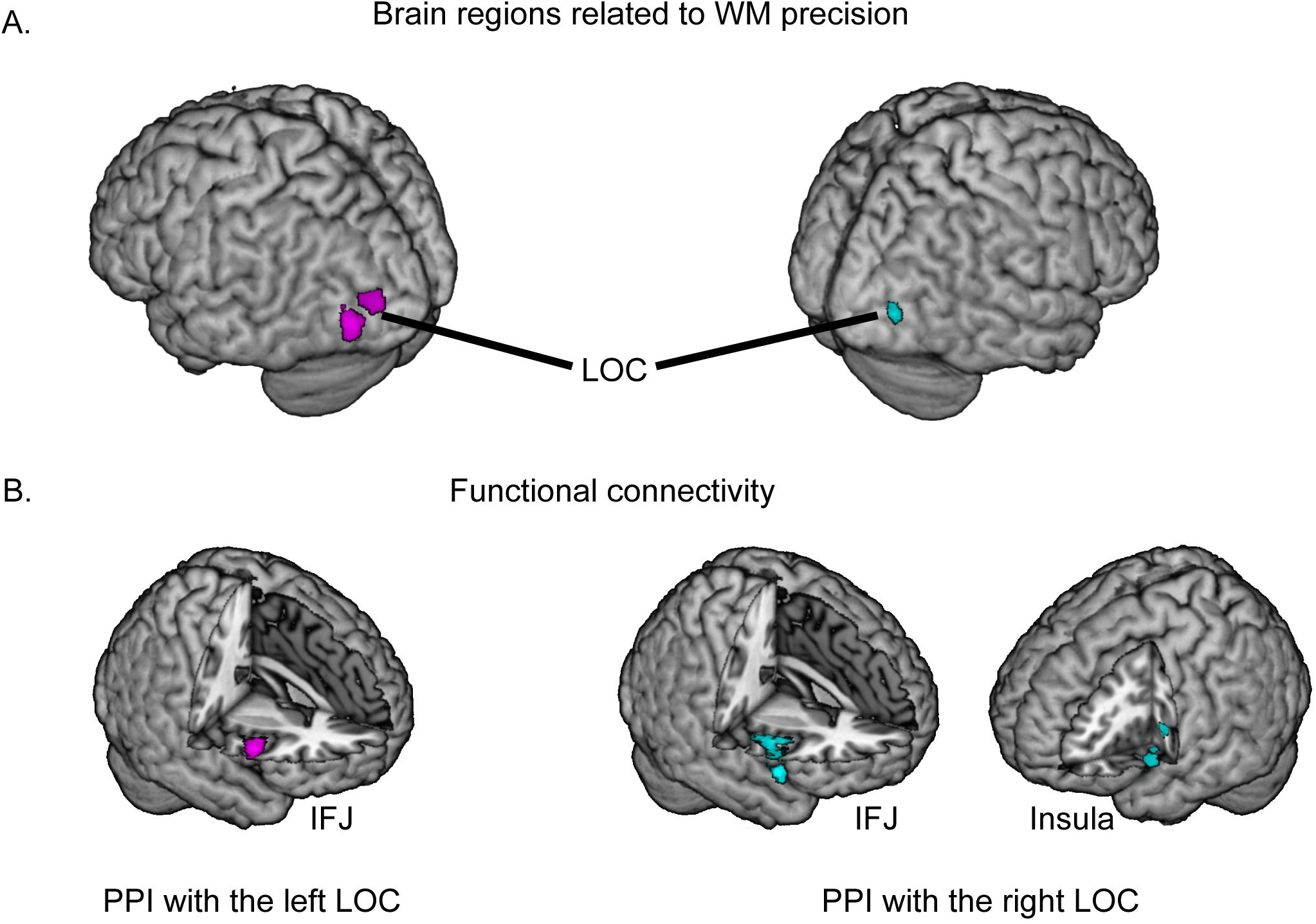
Parametric analysis and PPI results. A. Localization of brain regions reflecting precision. We included the absolute value of the response error observed on each trial in a GLM analysis (see Methods). Three clusters in lateral occipital cortex (two in left hemisphere and one in right hemisphere) exhibit significant negative beta weights on the response error regressors, indicating that activity levels in these regions are positively correlated with memory precision. B. Functional connectivity of precision-related regions and other parts of the brain. Two 5-mm–radian sphere ROIs centered at peak coordinates of clusters in A in each hemisphere respectively were set as seeds in PPI analysis. Both seeds exhibited stronger functional connectivity to right inferior frontal junction (IFJ) in high working memory load trials (load 4 & 6) compared with low working memory load trials (load 1 &2). Right lateral occipital cortex seed was also connected with left insula at higher working memory load.

However, even though the “precision ROIs” exhibited response profiles that tracked the maintenance of VWM precision, it is unlikely that this kind of process is completely independent of the WM load effect. Here, we performed a PPI analysis to see if the ROIs are functionally connected with other regions in the brain that are sensitive to VWM load. Note that since the two clusters in the left hemisphere are so close to each other, we combined them into one ROI and set the PPI seed at the center of the peak voxel in this combined ROI. Results in Figure 4B shows that both ROIs exhibited significantly higher connectivity with the right inferior frontal junction when the load increased (IFJ, high load > low load, l_LOC: t_(12)_ = 8.528, 309 voxels; r_LOC: t_(12)_ = 5.982, 198 voxels). In addition, the right LOC also showed differential functional connectivity with the left insula between the high and low WM load (t_(12)_ = 4.906, *p*_FWE_ < 0.05, 72 voxels).

### Relating brain activity and response precision

To test whether the areas activated during the WM task really trace the WM precision at the individual level, we applied a correlation method (Bland and Altman 1995) which focused on the within-subject changes across load by using an ANCOVA to eliminate the between-subject differences in BOLD signal. The results showed that the average beta values in all “precision ROIs” during the delay period across the four memory loads were correlated to the WM precision estimated from the mixture model using the data from experiment 1 (r = 0.732, *p* < 0.001; Fig. 5).

**Figure 5.**
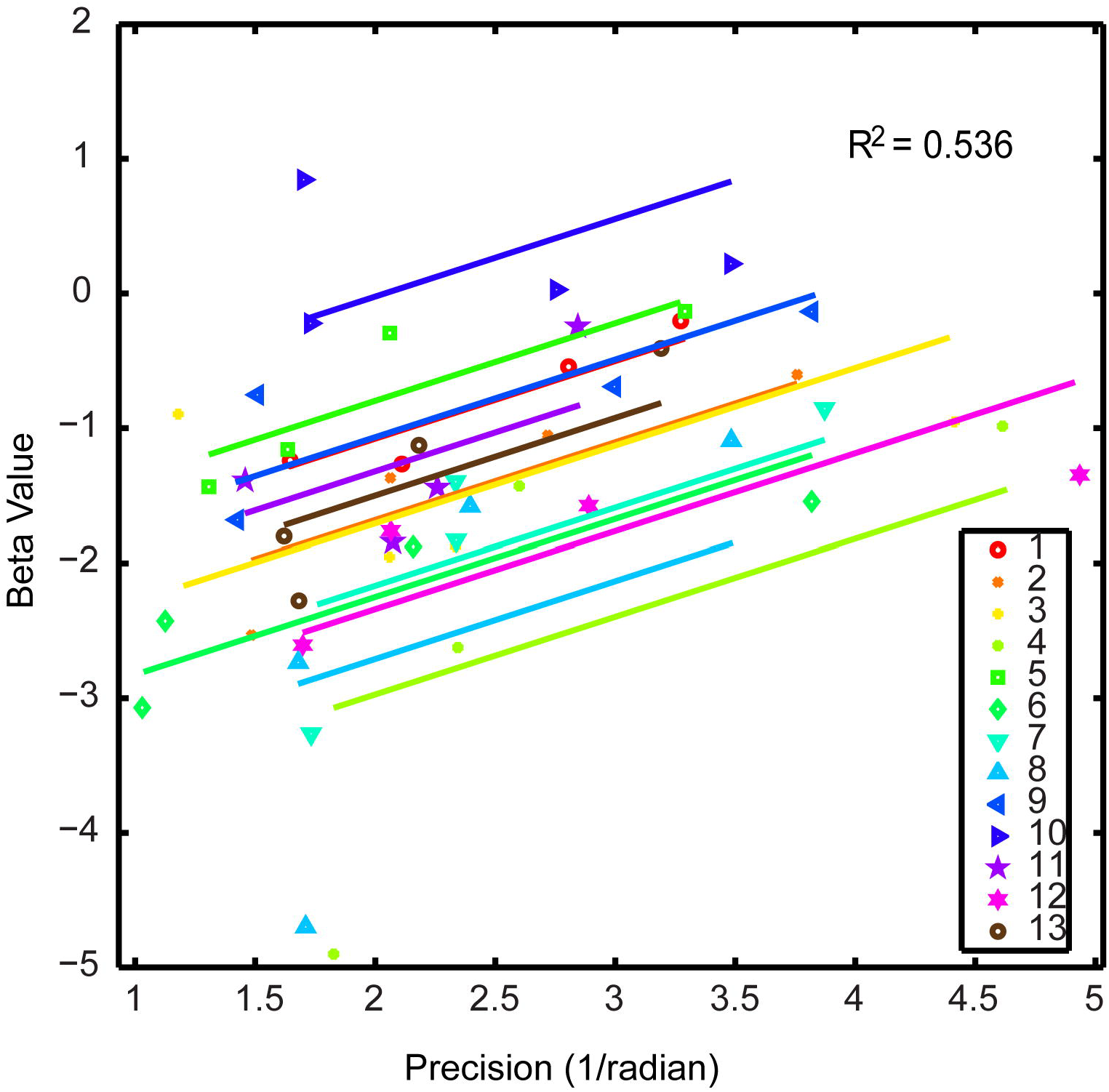
The relationship between brain activities in precision-related brain regions and behavioral performance. There are four dots for each subject, where each dot indicates for one WM load level. Resutls showed that changes in BOLD signal across WM loads were significantly correlated with changes in working memory precision returned from the mixture model (Bays et al. 2009) in the behavioral experiment outside MRI scanner at individual level. Data were modeled for each subject and fit with parallel lines with ANCOVA according to the methods of Bland and Altman (1995) (Bland and Altman 1995).

## Discussion

In order to characterize the neural mechanisms associated with the precision of VWM, in the current study, an orientation recall task was performed inside and outside an fMRI scanner under different VWM loads. Consistent with previous findings, behavioral results revealed that the precision decreased and the guessing rate increased when WM load was increased (Zhang and Luck 2008; Bays et al. 2009; van den Berg et al. 2012). Interestingly, BOLD signal in LOC could trace the precision of VWM trial-by-trial inside the MRI scanner, and were further correlated with the precision of VWM calculated from the behavioral performance outside the scanner at the individual level by a mixture model (Bays et al. 2009). Therefore, LOC is likely an important candidate for neural mechanisms underlying the precision of VWM. These results coincide with Xu and Chun’s previous findings, which indicate the delay activity in LOC co-vary with object complexity (Xu and Chun 2006). Meanwhile, our results did not contradict directly with previous findings where primary visual cortices (Harrison and Tong 2009; Sprague et al. 2014), parietal cortices (Bettencourt and Xu 2015; Ester et al. 2015) or frontal cortices (Ester et al. 2015) encode stimulus-specific mnemonic representations during VWM, as those studies all used multivariate methods to decode mnemonic objects using activity in those brain regions. However, none of those decoded patterns linked to the trial-by-trial performance, which was established in the present study. We used the behavioral response error as a parametric regressor to model a univariate GLM and only LOC exhibited a significant negative correlation with the error, which implicates the sensitivity of LOC activity to the precision of VWM. It is urged for future studies to link the decoding accuracy from the neural activity during the periods of sensory encoding and memory maintenance with the trial-wise precision from the behavioral data.

LOC, which is located in the middle level of visual hierarchies, is appropriate for connecting sensory-specific representations and memory-specific representations. Importantly, the connectivity between the bilateral LOCs and the right IFJ is stronger when the task load increases, which provides further evidence for a distributed network to represent different aspects of targets in VWM (Ku et al. 2015) and to fulfill goal-directed actions. This functional connectivity could be either a bottom-up information transfer or a top-down modulation of sensory cortical activity. Here, we speculate it is more likely a top-down control from the prefrontal cortex since previous research has provided causal evidence that top-down signals originating from the prefrontal cortex modulate sensory cortical activities (Ruff et al. 2006; Feredoes et al. 2011). Furthermore, applying inhibitory repetitive TMS to the right IFJ decreases neural suppression (i.e., increases activity) in sensory cortices to irrelevant stimuli and results in an increased reliance on LOC to uphold VWM performance (Zanto et al. 2013). Therefore, the current finding provides further evidence for the interaction between the prefrontal cortex and sensory cortices (Ku et al. 2015) and supports the distributed nature of VWM (Christophel et al. 2017).

It is notable that seeds within the bilateral LOC both increased the connectivity to the right IFJ with increasing VWM load. This hemispheric asymmetry within the prefrontal cortex is consistent with previous findings that revealed the involvement of the right prefrontal cortex more frequently than the left prefrontal cortex during visuospatial WM tasks (Postle and D’Esposito 2000; Gazzaley et al. 2004; Zanto et al. 2011). Addionally, when the right LOC is set as a seed, connectivity to the left insula is observed as well. The left insula and the right IFJ in our PPI results echoed a meta-analysis study showing these two regions’ critical function in intrusion resistance (Nee and Brown 2013), which may help explain why functional connectivity is stronger in the high-load condition – possibly to protect the VWM contents from internal interference. Moreover, insula has also been suggested to be involved in sustained attention during the maintenance of WM (Dosenbach et al. 2008) and recruited to support performance in high-demanding tasks (Derrfuss et al. 2005; Minzenberg et al. 2009). Previous research also showed evidence that the insula is involved in multimodal WM tasks including visual (Pouthas et al. 2005), auditory (Bamiou et al. 2003; Arnott 2005) and tactile (Sörös et al. 2007) stimuli, which further indicates its role in encoding multiple sourses of sensory information (Silverstein et al. 2010) and goal-directed actions.

Parietal cortices, especially the superior IPS, is critical in maintaining robust WM representation against distractors (Behrmann et al. 2004; Bettencourt and Xu 2015), and possibly in controlling the trial-by-trial variability of WM precision (Galeano Weber et al. 2016). While the current study did not observe a direct correlation between parietal activity and the trial-by-trial response error, IPS is more critical when more mnemonic information is needed to be maintained in WM (i.e., to represent multiple memorandums) (Xu 2017). Indeed, in the present study, the superior and inferior IPS is activated when the load of WM increases, which coincide with previous findings (Todd and Marois 2004; Barch et al. 2013).

In conclusion, the current study provides evidence that the LOC plays an important role in maintaining detailed visual representations in VWM, and increased functional connectivity between LOC and prefrontal areas such as the right IFJ and the left insula help fulfill goal-directed VWM processes under increased task demands (i.e., when the VWM load is higher).

## Acknowledgement

We thank Szechai Kowk, Qing Cai for valuable suggestions about data analysis, Wen Fang for suggestions about analytical programming, and Yong-Di Zhou for helpful comments on the manuscript. This work was supported by the National Key Fundamental Research Program (2013CB329501), the Major Program of Science and Technology Commission Shanghai Municipal (17JC1404100), the Shanghai Pujiang Talents Plan Project (16PJC022), and the NYU-ECNU Institute of Brain and Cognitive Science at NYU.

